# Modulation of sperm capacitation enhances blastocyst hatching in bovine in vitro fertilization

**DOI:** 10.64898/2026.03.18.712589

**Authors:** Olinda Briski, Fernanda Fagali Franchi, Ernesto Piga, Federica Franciosi, Sai Kamal Nag Bonumallu, Carolina Baro Graf, Valentina Lodde, Alberto Maria Luciano, Dario Krapf

## Abstract

In vitro fertilization (IVF) is key for genetic improvement programs in bovine. However, embryos produced through IVF have lower developmental competence than those produced under *in vivo* conditions. Conventional sperm preparation for IVF typically relies on heparin for sperm capacitation but fails to replicate the finely tuned molecular environment of the oviduct, resulting in compromised embryonic competence. Here, we evaluated the effect of *HyperBull*, a novel capacitation technology, on bovine IVF outcomes using unsorted cryopreserved semen. In a split-sample design, 528 cumulus–oocyte complexes were co-incubated with either control or *HyperBull* capacitated spermatozoa from the same bull. While overall blastocyst rates were not significantly different between groups (34.21% *HyperBull* vs. 28.63% control, p=0.148), the proportion of hatched embryos was significantly higher in the *HyperBull* group (15.82% vs. 9.13%, p=0.016). These findings suggest that modulating capacitation signals prior to insemination enhances embryonic developmental competence, thereby improving readiness for implantation. *HyperBull* may thus represent a valuable tool to increase the efficiency of IVF programs.

## INTRODUCTION

Embryo production in bovine species largely relies on *in vitro* fertilization (IVF), a crucial step in livestock breeding. Despite advances, the efficient production of viable embryos through IVF remains a significant challenge in the field. The proportion of produced embryos suitable for transfer generally ranges between 20% and 40%, depending on the in vitro culture system employed (Ferré et al., 2020). The limited yield of transferable embryos *in vitro* is not primarily due to failures in oocyte maturation or fertilization, as nuclear maturation and fertilization rates are typically above 70% (Lonergan and Fair, 2014). One of the main issues is in suboptimal conditions during *in vitro* sperm capacitation, resulting in inadequate development of many fertilized embryos during culture(Osycka-Salut et al., 2024a; Gómez-Elías et al., 2025). We have previously shown that improving spermatozoa capacitation *in vitro* yielded higher blastocyst rates in bovine (Osycka-Salut et al., 2024a). Given the variability in sperm quality across bulls and the fact that bulls are generally not selected for IVF performance, improving *in vitro* fertilization outcomes may benefit from enhanced sperm selection and/or capacitation. For instance, methods based on rheotaxis have increased the proportion of embryos reaching the blastocyst stage compared to conventional centrifugation isolation. Another promising avenue is mimicking the natural interactions between spermatozoa and the oviduct. Spermatozoa binding to oviductal epithelial cells promotes their survival (Pollard et al., 1991a), and studies in rabbits and pigs suggest that the oviductal isthmus may have a role in reducing polyspermy (Mahé et al., 2021a). Spermatozoa capacitation *in vivo* involves complex modifications mediated by the oviduct (Mahé et al., 2021a; Delgado-Bermúdez et al., 2022a). *In vitro*, capacitation is typically induced by heparin(Parrish, 2014), which may not fully replicate the oviductal influences on spermatozoa’s function. Delayed or incomplete capacitation could result in fertilization by defective spermatozoa and reduce embryo competence (Koyama et al., 2014). Significantly, spermatozoa differ in their capacity to produce embryos with high developmental potential (Hansen et al., 2010; Vallet-Buisan et al., 2023; Yaghoobi et al., 2024). Fertilization with defective spermatozoa may be more common *in vitro* than *in vivo*. While spermatozoa concentrations *in vitro* typically approximate 1 × 10^6/mL, the number of spermatozoa reaching the fertilization site *in vivo* is considerably lower, often in the hundreds or even close to 1:1 in mice (La Spina et al., 2016). The natural capacitation process within the female reproductive tract likely endows spermatozoa with superior fertilizing ability and embryonic support (Miller, 2024).

This highlights an urgent need to reconsider and improve *in vitro* spermatozoa capacitation. Here, we show how modulating capacitation signals in the spermatozoa improves blastocyst quality, resulting in an increase in hatched embryos after conventional *in vitro* fertilization.

## METHODS

All chemicals and reagents used in this study were purchased from Merck Sigma Aldrich, Italy, except those specifically mentioned. Disposable sterile plasticware was purchased from SARSTEDT Srl, Italy (SARSTEDT Green line for suspension cells) and Thermo Fisher Scientific Inc, Germany (NUNC IVF Line and Sterilin™). All the procedures were conducted at room temperature (26°C) unless otherwise specified.

### Animal Welfare

Bovine ovaries were collected post-mortem from healthy cows subjected to routine veterinary inspection at a licensed slaughterhouse (Inalca SpA), in compliance with European and national legislation (EC No. 1069/2009; EC No. 1099/2009). No animals were killed specifically for this study. Oocytes were used for in vitro fertilization with commercial bull semen, and embryos were cultured *in vitro* up to the blastocyst stage. In accordance with Directive 2010/63/EU, preimplantation embryos do not fall under the scope of animal experimentation regulations; therefore, no ethical approval was required.

### Cumulus-Oocyte Complexes Collection and In Vitro Maturation

Holstein Friesian bovine ovaries were recovered at a local abattoir (IT 2270M CE; Inalca S.p.A., Ospedaletto Lodigiano, LO, Italy) from pubertal dairy cows (4-8 years old) subjected to routine veterinary inspection and according to the specific health requirements. Only animals with both ovaries with more than 10 mid-antral follicles (2-8 mm) visible on the ovarian surface were considered (Modina et al., 2014). Ovaries were transported in saline solution at 28-32 °C. As previously described (Luciano et al., 2005), Cumulus–oocyte complexes (COCs) were retrieved from medium antral follicles with a 16-gauge needle mounted on an aspiration pump (COOK-IVF, Brisbane, QLD, Australia) (Luciano et al., 2005). COCs were washed in TCM-199 supplemented with HEPES buffer 20 mM, 1790 U/L heparin, and 0.4% of bovine serum albumin (BSA) (HM199) and examined under a stereomicroscope. Only COCs medium-brown in color with five or more complete layers of cumulus cells with oocytes with finely granulated homogeneous ooplasm were used.

COCs were washed twice in HM199, then cultured in groups of 30-35 in 500 µl of IVM medium (M199 with Earle′s salts supplemented with 25 mM sodium bicarbonate, 2 mM GlutaMAX™, 0.4% fatty acid free bovine serum albumin, 0.2 mM sodium pyruvate, 0.1 mM cysteamine, 50 µg/ml of kanamycin, 0.1 IU/ml of r-hFSH) for 24 h, in four-well dishes at 38.5°C under 5% CO_2_ in humidified air (Luciano et al., 2005). A total of 528 COCs were used across three biological replicates (*HyperBull*: n = 266; control: n = 262).

### In Vitro Fertilization and Embryo Culture

In vitro fertilization (IVF) and in vitro culture (IVC) were performed as previously described, with slight modifications (Tessaro et al., 2015). Straws of cryopreserved Holstein bull spermatozoa (CIZ, S. Miniato, Pisa, Italy) were selected using a 45-90% Percoll–TALP discontinuous gradient and subsequently centrifuged at 700 × g for 30 minutes. The supernatant was removed, leaving approximately 200 µL of pellet, and then 350 µL of TALP medium was added to the pellet (final spermatozoa solution ∼ 550 µL, See Fig. 1 for a graphical protocol). 250 µL of the spermatozoa suspension was placed in each of to two tubes. For *HyperBull* treatment, specific reagents were added, while for the control, the same volume of *HyperBull* was replaced with TALP. Then, 3 ml of TALP was added to both tubes (*HyperBull* and control), and centrifuged for 10 min at 400 × g. Following centrifugation, the supernatants were removed, and the spermatozoa were counted using a Neubauer chamber. The spermatozoa were added to the four wells containing TALP-IVF supplemented with PHE (1 mM hypotaurine, 2 mM penicillamine, and 250 mM epinephrine), and matured COCs to reach a final concentration of 750,000 sperm/ml. A total of 70–90 COCs were inseminated per group, in 3 biological replicates, for both the *HyperBull* and control conditions. After 18h of IVF, presumptive zygotes were cultured as previously reported (see (Luciano et al., 2005)). Briefly, residual cumulus cells and spermatozoa were removed by vortexing for 90 sec in 500 µL of synthetic oviduct fluid buffered with 10 mM HEPES and 5 mM NaHCO_3_ (SOF wash), rinsed twice, and then transferred into SOF embryo culture medium (Luciano et al., 2005). The embryo culture medium (SOF-IVC) was synthetic oviduct fluid buffered with 25 mM NaHCO_3_ and supplemented with MEM essential and nonessential amino acids, 0.72 mM sodium pyruvate, 2.74 mM myo-inositol, 0.34 mM sodium citrate, and 5% calf serum (CS). Incubation was performed at 38.5 °C with a humidified gas mixture composed of 5% CO_2_, 5% O_2_, and 90% N_2_. On day 8, blastocyst rates and morphology were assessed, as previously described (Franciosi et al., 2014). Briefly, blastocysts in which the blastocoel had just begun to form, and the cell types were not distinguishable, were classified as not expanded. Blastocysts were classified as expanded if the blastocoel was fully formed and the trophectoderm and inner cell mass were clearly distinguishable but still contained within a thinned zona pellucida. Hatched blastocysts were those outside the zona pellucida.

**Figure 1.**
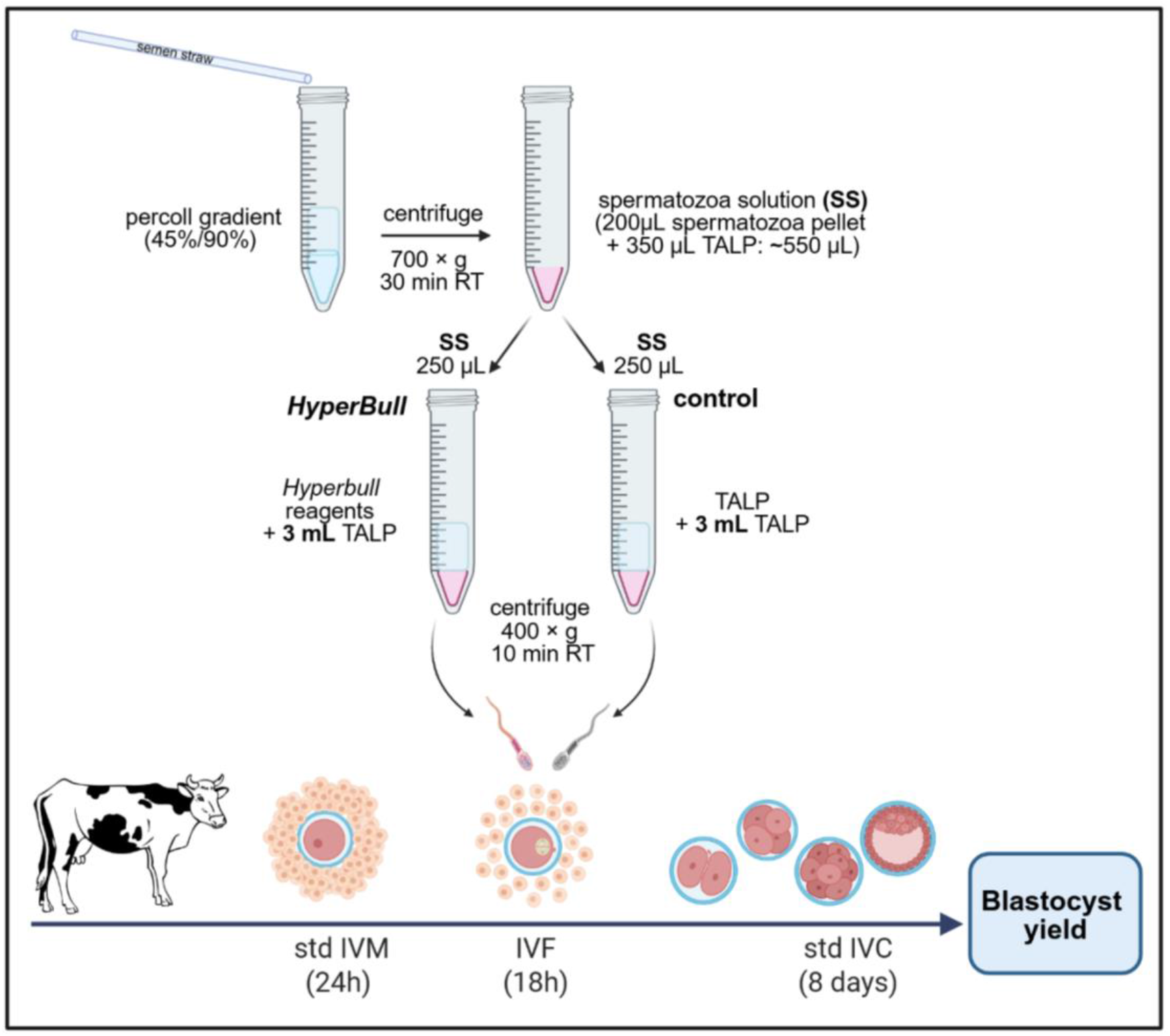
Experimental design. Slaughterhouse-derived COCs were selected and in vitro matured for 24 h. Matured COCs were co-incubated with spermatozoa pre-processed using either the standard protocol (control) or *HyperBull* for 18 h. A total of 528 COCs were inseminated across three biological replicates for both *HyperBull* and control conditions (n = 266 and 262, respectively). The presumptive zygotes were freed from remaining cumulus cells and cultured for 8 days under standard in vitro culture conditions. On day 8, blastocyst development and morphology were evaluated to assess IVP outcomes.

### Statistical Analysis

Blastocyst rate was calculated as the number of blastocysts per total cultured oocytes, and hatching rate as the number of hatched blastocysts per total blastocysts. Blastocyst and hatching rates were expressed as percentages and analyzed as paired data, within each replicate block. The normality of paired differences was evaluated using the Shapiro– Wilk test, which showed no significant deviation from normality for either endpoint. Therefore, paired Student’s t-tests were used to assess treatment effects.

Statistical significance was set at p < 0.05. Analyses were performed using GraphPad.

## RESULTS

### *HyperBull* improves hatching of in vitro produced bovine embryos

The test comparing *HyperBull* performance to standard protocols in terms of blastocyst developmental rates shows that *HyperBull* yielded a higher number of blastocysts than the control (34.21% vs. 28.63%, respectively; Table 1), although this difference was not statistically significant. A binomial logistic regression analysis comparing blastocyst rates in the control and *HyperBull* groups showed a treatment coefficient of +0.26, indicating a trend toward a higher blastocyst ratio. Most notably, the proportion of hatched blastocysts in the *HyperBull* group was significantly higher compared to the control (15.82% vs. 9.13%, p=0.016).

**Table I.**
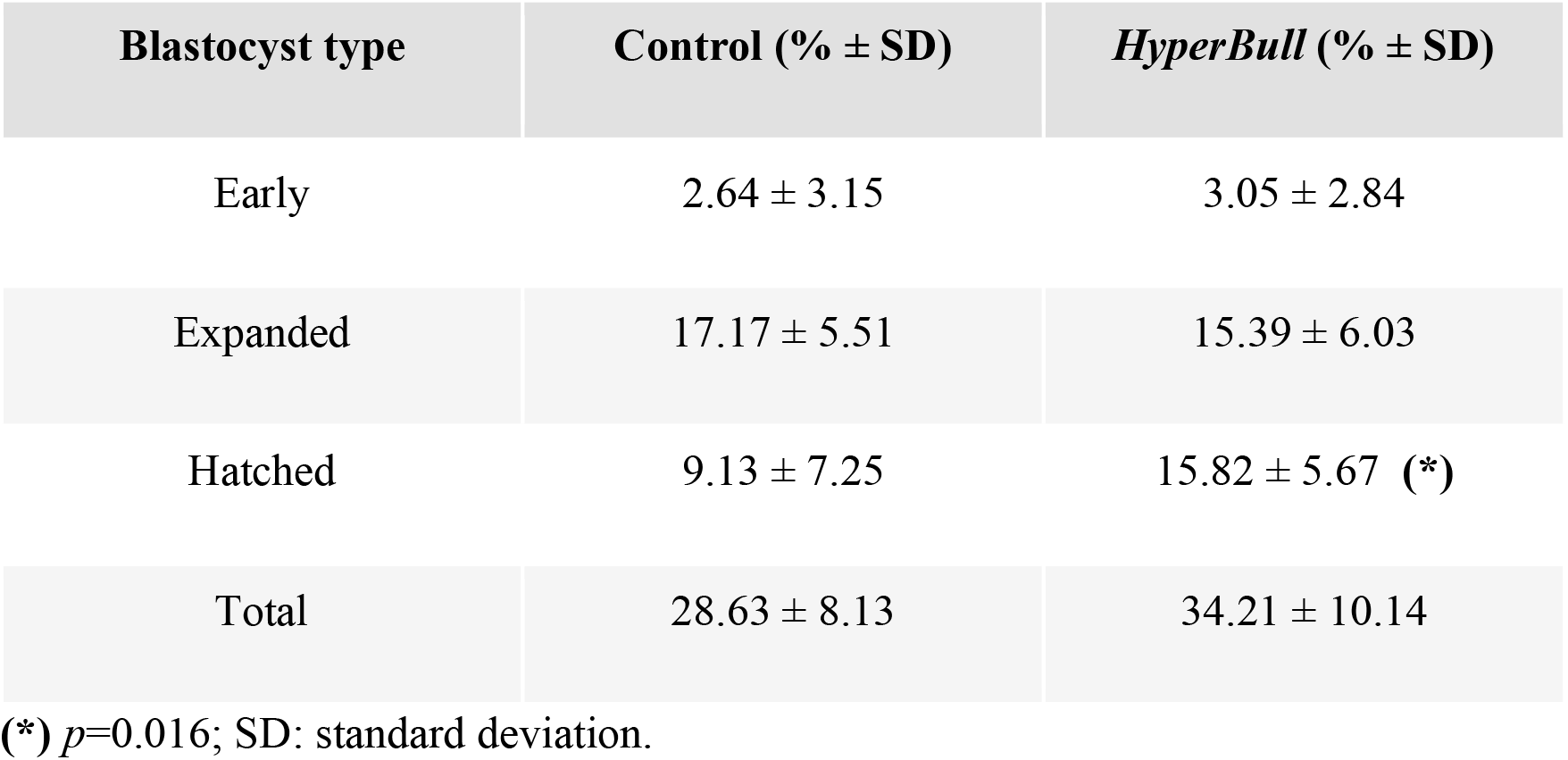
Effect of *HyperBull* spermatozoa treatment on blastocysts yield. Three biological replicates, with at least two technical replicates per procedure (8 technical replicates in total), were performed, with a total of 266 COCs assigned to *the HyperBull* treatment and 262 COCs assigned to the control.

| **Blastocyst type** | **Control (% ± SD)** | ***HyperBull* (% ± SD)** |
| --- | --- | --- |
| Early | 2.64 ± 3.15 | 3.05 ± 2.84 |
| Expanded | 17.17 ± 5.51 | 15.39 ± 6.03 |
| Hatched | 9.13 ± 7.25 | 15.82 ± 5.67 (*) |
| Total | 28.63 ± 8.13 | 34.21 ± 10.14 |

(\*) $p=0.016$ ; SD: standard deviation.

## DISCUSSION

The present study suggests that targeted manipulation of bovine sperm capacitation with *HyperBull* enhances the embryo quality by increasing the proportion of hatched blastocysts following IVF. Although overall blastocyst rates showed an upward trend compared to controls, the marked increase in hatching rates suggests that *HyperBull* confers functional benefits that appear later in embryonic development. This finding aligns with the hypothesis that optimizing capacitation signals before fertilization improves the developmental competence of resulting embryos, even when early cleavage and blastocyst formation rates remain relatively stable (Parrish et al., 1988; Rizos et al., 2002; Coy et al., 2012).

These results should be viewed in the context of the limitations of *in vitro* reproductive technologies. While the oocyte nuclear maturation and fertilization rates are generally high, the early embryo development is often suboptimal. As highlighted in previous studies (Rizos et al., 2002; Gad et al., 2011), embryos derived entirely *in vitro* exhibit inherent deficits in competence, partially due to suboptimal sperm capacitation conditions (Navarrete et al., 2019). Wheater *in vivo*, the bovine capacitation in the oviduct is a finely tuned sequence of molecular events, involving gradual membrane remodeling, regulation of intracellular ion homeostasis, and modulation by oviductal secretions (Mahé et al., 2021b; Delgado-Bermúdez et al., 2022b); the standard *in vitro* capacitation protocols most often relying on heparin supplementation, do not replicating the spatial and temporal signaling gradients present in *vivo*, potentially leading to incomplete capacitation.

Capacitation influences not only the ability of the spermatozoon to penetrate the oocyte but also the integrity and remodeling of the paternal genome, centrosome function, and the activation of embryonic transcription (Simon et al., 2014; Lismer and Kimmins, 2023; Xie et al., 2023). Indeed, the alternative procedures for spermatozoa capacitation in bovine and murine models lead to increased embryo development (Tateno et al., 2013; Osycka-Salut et al., 2024a). In those studies, manipulation of capacitation signaling not only improved fertilization efficiency but also exerted synergistic effects on blastocyst production when combined with conventional inducers such as heparin. *HyperBull* appears to operate on a similar principle, modulating capacitation pathways beyond the conventional heparin-induced mechanism resulting in the improvement of blastocysts hatching rates.

Comparable improvements in embryonic developmental competence have been reported with alternative sperm-selection strategies, with approaches such as rheotaxis-based selection (Yaghoobi et al., 2024) or mimicking oviductal epithelial binding (Pollard et al., 1991b) generating embryos of higher developmental potential than those derived from standard density-gradient or swim-up techniques *HyperBull*’s effects may be complementary to these approaches, creating an opportunity to combine both strategies within a single protocol to further enhance embryo yield and overall reproductive efficiency.

From an applied perspective, the *HyperBull* shows particular a promise for improving outcomes when using challenging sperm sources, such as sex-sorted or cryopreserved semen, which often have reduced motility and lower IVF success rates (Suh et al., 2005). By enhancing hatching rates, this approach could benefit commercial bovine breeding programs where maximizing genetic gain is essential. Overall, these findings reinforce that optimizing sperm capacitation prior to insemination can enhance embryo quality, positioning *HyperBull* as a valuable addition to bovine assisted reproduction systems. Future research should evaluate its compatibility with advanced sperm-selection methods, determine its efficacy across diverse bull populations, and critically assess whether the observed *in vitro* gains translate into higher pregnancy rates and improved calf outcomes in vivo.

## CONCLUSION

Performing sperm capacitation using HyperBull significantly increased blastocyst hatching rates in bovine IVF without altering overall blastocyst yield. These results indicate that optimizing capacitation signals prior to fertilization enhances embryo quality and developmental competence. HyperBull may therefore represent a valuable tool to improve the efficiency of bovine IVF systems, particularly when using cryopreserved semen.

## Author contributions

**Olinda Briski:** Methodology, Writing – review and editing, Investigation; **Fernanda Fagali Franchi:** Methodology, Writing - review and editing, Formal análisis; **Ernesto Piga**: Methodology; **Federica Francios:** methodology; **Sai Kamal Nag Bonumallu:** Investigation; **Carolina Baro Graf**: Methodology, Project administration; **Valentina Lodde:** Data curation, Formal análisis, Methodology, Writing – review and editing, Supervision, Validation; **Alberto Maria Luciano:** Data curation, Formal análisis, methodology, Writing – review and editing, Supervision; Validation; **Dario Krapf:** Data curation, Formal análisis, Methodology, Writing – review and editing, Supervision, Conceptualization

